# Tracking Conditioned Fear in Pair-Housed Mice Using Deep Learning and Real-Time Cue Delivery

**DOI:** 10.1101/2025.05.10.653260

**Authors:** Hannah C. Smith, Zhe Yu, Patrick Yazigi, Benjamin Turley, Adam Swiercz, Jeanie Park, Paul J. Marvar

**Affiliations:** Departments of Neuroscience and Pharmacology & Physiology, George Washington University, Washington, DC 20052; School of Medicine, Georgetown University, Washington, DC 20057; National Institute of Mental Health, National Institutes of Health, Bethesda, MD 20892; Department of Medicine, Emory University, Atlanta, GA 30322

## Abstract

Post-traumatic stress disorder (PTSD) is a complex and prevalent neuropsychiatric condition that arises in response to exposure to a traumatic event. A common diagnostic criterion for PTSD includes heightened physiological reactivity to trauma-related sensory cues, in safe or familiar environments. Understanding complex PTSD criteria requires new pre-clinical paradigms and technologies that integrate sensory physiology (e.g., auditory, visual, olfactory) with behavior. Here we present a novel Pavlovian-based paradigm using an open-source software plus deep learning-based pose estimation to investigate the effects of a recurrent conditioned stimulus (CS) on fear behaviors in pair-housed mice within the home cage.

Simultaneous home cage video recording and analysis of CS-evoked freezing behaviors were performed using a deep learning model, with consideration for light-dark circadian cycles. Fear-conditioned dyad mice exhibited high CS-evoked freezing, with evidence of extinction learning (characterized by low freezing) during the mid-phase of the 2-week paradigm. Females exhibited reduced CS-evoked home cage freezing compared to males with circadian differences between the light (low freezing) and dark (high freezing) periods. Following the 2-week paradigm, fear-conditioned mice, compared to controls, exhibited heightened context-dependent freezing, while males but not females showed heightened startle reactivity. Taken together, these results demonstrate a novel software application for examining conditioned defensive and fear behaviors over time in mouse dyads within an ethologically relevant environment. Future applications could be used for more integrative analysis and understanding of neural circuits and heightened sensory threat reactivity, potentially improving the understanding and treatment of PTSD.

## INTRODUCTION

PTSD is a prevalent heterogeneous psychiatric disorder with notable sex differences and physical health comorbidities(Bryant, 2019; Fonkoue et al., 2020; Ressler et al., 2022). As outlined in the Clinician-Administered PTSD Scale for DSM-5 (CAPS-5), PTSD is characterized by impaired sensory processing, which can manifest as intrusive symptoms like recurrent flashbacks and the overgeneralization of trauma-related sensory cues(Fleming et al., 2024).

These sensory disturbances can contribute to a range of symptoms, including hypervigilance, avoidance behaviors, and exaggerated physiological and autonomic responses, such as increased heart rate and blood pressure (Hicks et al., 2024; Lee et al., 2022; Putica and Agathos, 2024). However, these underlying mechanisms and manifestations of PTSD clinical symptoms remain unclear. To address this, there is a growing need for improved pre-clinical research and technologies that integrate physiological and behavioral applications with multi- dimensional analysis in ethologically relevant environments, such as the home cage (Hegoburu et al., 2024; Kahnau et al., 2023; Signoret-Genest et al., 2023; Tanas et al., 2022).

Pavlovian fear conditioning, widely used in pre-clinical and clinical PTSD research, has been instrumental in identifying neural signatures linked to fear and maladaptive fear memory in PTSD (Bowers and Ressler, 2015; Dunsmoor et al., 2022; Richter-Levin et al., 2019) In addition, our laboratory(Swiercz et al., 2018; Turley et al., 2021) and others (Stiedl et al., 2009, 2004; Tovote et al., 2005), have demonstrated the utility of integrating behavioral and cardio-autonomic (e.g., heart rate and blood pressure) analysis of conditioned fear and defensive behaviors, with real-time wireless telemetry recordings, with important implications for translating findings to humans (Seligowski et al., 2020). Although Pavlovian behavioral paradigms are arguably one of the most widely used pre-clinical PTSD paradigms, these paradigms are often limited to the study of short-term phasic fear or defensive behavioral responses (ie., escape or freezing) of one animal in a specified behavioral arena context. Moreover, there is limited understanding of the neurobiology of conditioned fear-related behaviors in ethologically relevant social contexts (Hegoburu et al., 2024; Ito et al., 2023) and how impaired sensory cues (e.g., visual, auditory, olfactory) affect circadian rhythms (Hartsock et al., 2023) and reward-seeking or anxiety-like behaviors (Chu et al., 2024), in these contexts.

Building on this gap, we recently developed a customizable closed-loop add-on software for Ponemah™ that considers the diurnal and resting cardiovascular state of a mouse while in its home cage, enabling remote delivery of a conditioned stimulus (e.g., auditory tone) based on a predefined cardiovascular state threshold(Turley et al., 2021). To further examine conditioned fear in ethologically relevant environments, this open-source software also has a feature for the remote programming of pseudorandomized sensory cues, delivered day or night within the home cage, a feature not previously tested. Therefore, the current study objective was to test this feature of the software called *Adaptive Cue Delivery* (ACD – GitHub <u>Link</u>) using a Pavlovian paradigm to assess the effects of intermittent, pseudorandomized auditory cue delivery to the home cage on fear behaviors in pair-housed mice across circadian cycles.

## MATERIALS and METHODS

### Animals

All experimental procedures were approved by the Institutional Care and Use Committee (IACUC) of the George Washington University and followed National Institutes of Health guidelines. The studies used 8–10-week-old male and female C57/BL/6 mice pair-housed mice in a temperature and humidity-controlled room on a 12hr light/dark cycle. Food and water were available ad libitum.

### Auditory cue-dependent Pavlovian fear conditioning

Pavlovian fear conditioning, as previously described by our lab(Hurt et al., n.d.; Marvar et al., 2014; Smith et al., 2024; Yu et al., 2019), was used to model associative fear by pairing an auditory cue with a light foot shock.

Mice were habituated to the fear conditioning chamber for two days (20min, 40min) before undergoing aversive associative conditioning in the light phase, during which mice received 20 conditioned stimulus (CS)/unconditioned stimulus (US) pairings using a 30s auditory cue (6kHz, 75dB, CS) co-terminating with a mild foot shock (1s, 0.6mA, US) at a variable inter-trial interval of 30s-1min. Male and female mice were fear conditioned (Colbuourn) individually using auditory fear conditioning, as previously described, and then tested 24 hours later as to pair-housed mice in a home cage.

### Home Cage CS Delivery with Adaptive Cue Delivery (ACD) program

Twenty-four hours after fear conditioning, the ACD software was pre-programmed to deliver 6 tones (6kHz, 75dB) over 24 hours, with 3 tones during the light (7am-7pm) and dark (7pm-7am) cycles to pair-housed mice. Inter-tone intervals were randomized, with a minimum of 1 hour between tones. Each cage had its own speaker and sound-attenuating panels to reduce external noise **(Figure 1A).** After each tone delivery day, a 24-hour break followed, repeating this cycle for 14 days **(Figure 1B),** 7 tone delivery days: experimental days 2, 4, 6, 8, 10, 12, 14). Cameras with day and night recording transmitted live video feed for tone-specific video extraction (Wisenet SNK- B73041BW, Hanwha Vision, Teaneck, NJ) and using a modified cage top with plexi-glass window (Clever Sys INC model# CSI-ENV-HC-ST).

**Fig 1.**
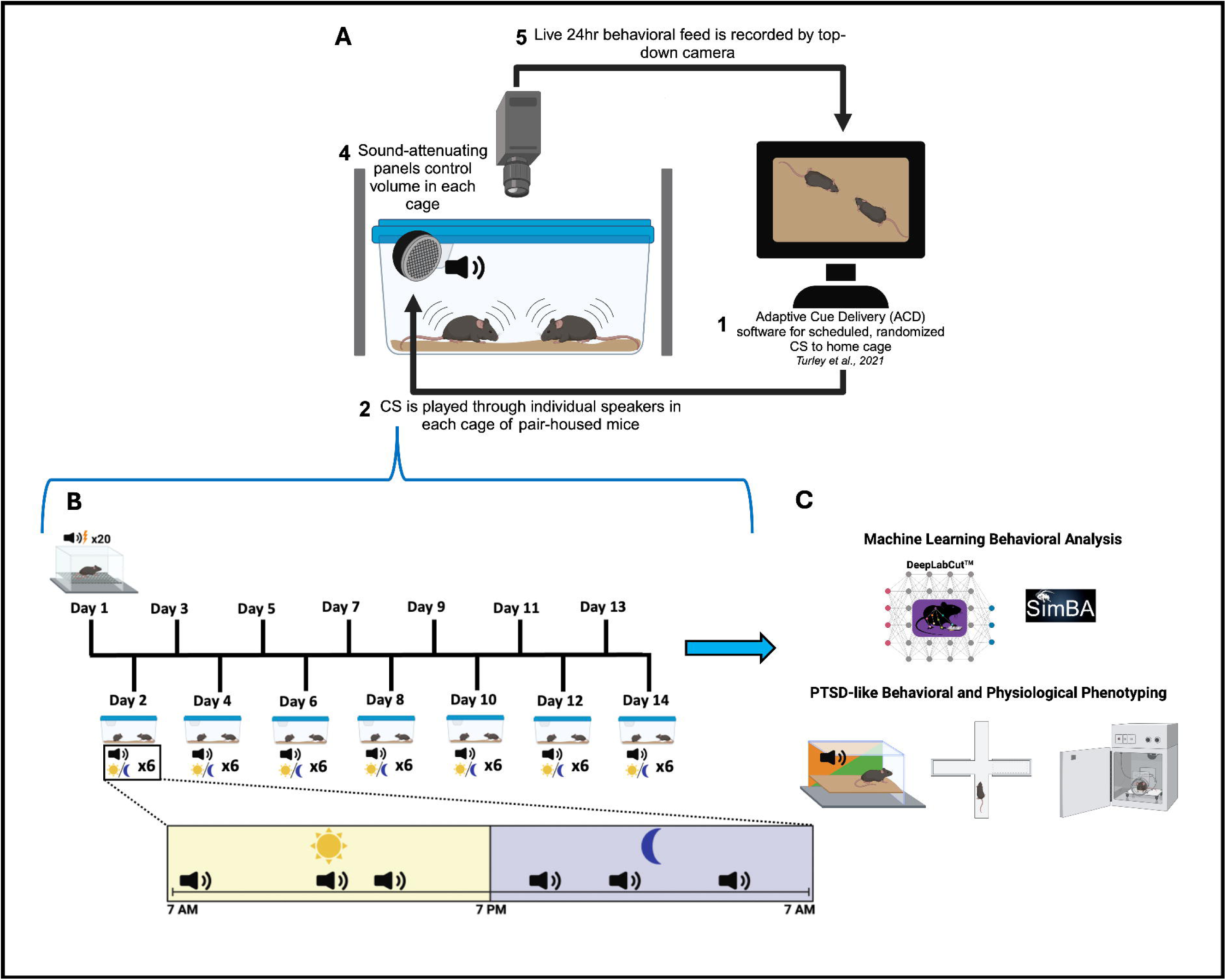
Protocol and experimental design. A. Integration of Adaptive Cue Delivery (ACD) with behavioral monitoring (1) ACD delivers conditioned auditory cues at random intervals over 24 hours. (2) Cues play through individual speakers in each cage of pair-housed mice (3) Cages are enclosed with sound-attenuating panels to standardize volume (4) Top-down cameras record freezing behavior continuously across light/dark cycles. B. Fear conditioning protocol and 14-day home-cage cue exposure. Conditioned auditory cues were delivered intermittently in the home cage over 14 days following fear conditioning. C. Machine learning analysis and behavioral phenotyping.

### Manual scoring of freezing behavior

For manual scoring of conditioned freezing behavior, two blinded observers independently assessed freezing behavior by reviewing extracted video recordings. As previously described, freezing was defined as the absence of visible movement, for at least 2 seconds, except that required for respiration (fluctuation in the volume of the thorax) (Fanselow and Bolles 1979) and was scored during each 30 CS – auditory cue of each cage during the first CS of each light/dark cycle (see supplemental videos). Unresponsive mice (e.g., sleeping) were not counted as freezing.

#### Deep learning-based pose estimation for tracking multi-animal behaviors

For automated freezing analysis in pair-housed mice, a deep neural network was trained using DeepLabCut (DLC) on behavior test videos, tracking six body points: nose, left ear, right ear, neck, body, and tail. After DLC tracking, the data was smoothed and missing points were filled using Simple Behavior Analysis (SimBA) software. A blinded experimenter then reviewed and corrected the data. Any misaligned data was excluded. Freezing was reliably detected based on low velocity of body points, with the left ear providing the most accurate detection. A 5 mm/s velocity threshold matched human annotations. This method effectively converts body point tracking into freezing data, enabling detection in dynamically changing environments (see supplemental file for expanded DLC methods).

### Assessing cue-evoked freezing in a novel context

To assess behavioral freezing of conditioned fear in a new context, on experimental day 15, following repeated intermittent CS reminders in the home cage, mice were placed in a novel context with altered visual, tactile, and olfactory cues and CS-evoked freezing was assessed. Following a 5min pre-CS interval, two CS (30s, 30s interval, 6kHz, 75dB) were given and freezing to each CS was recorded and calculated using Freezeframe 3.0 software (Actimetrics, Wilmette, IL).

### Assessing avoidance-like behaviors

To assess whether intermittent repeated CS delivery in the home cage impacted locomotion or avoidance-like behavior, on day 16, mice were placed into the elevated plus maze (EPM) as previously described by our lab(Marvar et al., 2014; Yu et al., 2023, 2019) . Briefly, mice were placed in the center of the apparatus facing the same closed arm and allowed to freely explore for 5min and open arm entries, and % time spent in the open arms of the maze were calculated using Anymaze software (Stoelting Co., Wood Dale, IL, USA).

#### Acoustic startle reflex (ASR) test

The next day following EPM testing, on experimental days 17-18, mice underwent ASR to evaluate mechanisms of alarm and heightened arousal. As previously described by our lab(Yu et al., 2024) mice were tested individually in four SR-Lab Startle units, each unit consisting of a transparent Plexiglas cylinder mounted on a platform attached to an accelerometer to measure millivolts (mV) startle amplitude (SR-Lab Startle Response System, San Diego Instruments, San Diego, CA). Each test session began by placing the mice into a Plexiglas cylinder for a 5 min acclimation, during which 70 dB background noise was maintained. After acclimation, each mouse was presented with 16 120 dB white noise startle bursts at random intervals over a 20-minute session. Startle responses were measured using SR-Lab software, which converts the millivolt signal from the sensor into a physical measurement, with millivolts proportional to the intensity of the startle response (force or movement).

### Fear potentiated startle (FPS) testing

Following ASR, on Day 18, mice underwent modified FPS testing(Smith et al., 2011), which consisted of a 5min acclimation period followed by 10 habituation trials of 40ms 110dB white noise bursts. After the leader trials, mice received a combination of 15 uncued trials (40ms, 110dB) identical to the leader trials, 15 cued trials (30s, 6kHz, 75dB tone co-terminating with 40ms, 110dB white noise burst), and 15 no-stimulation trials (30s of silence) presented in random order (Jones et al., 2005; Olivera-Pasilio and Dabrowska, 2023). Each trial was followed by a 30s inter-trial interval. FPS allows for both cued and uncued startle within one testing session, resulting in raw startle amplitude under several different conditions (habituation startle, uncued startle, cued startle) that allow for calculation of percent fear-potentiated startle exhibited during the test (Ayers et al., 2011; Moaddab and Dabrowska, 2017)

### Data presentation and statistical analysis

Data analysis was performed using GraphPad Prism 10.0 (GraphPad Software, CA), with outliers identified via the ROUT test (Q=1%). Mean differences between groups were assessed using one- and two-way ANOVA, with Bonferroni post-hoc analysis and Tukey’s multiple comparison tests. Statistical significance was set at p<0.05. Complete statistical results are provided in the Supplement Table.

## RESULTS

### CS-evoked freezing behavior in pair-housed mice within the home cage

Following Pavlovian auditory fear conditioning, CS (tone)-US (foot shock) pairings or CS-only mice were returned to their home cage. The ACD software, integrated with a video system, was set up to deliver six randomized CSs (three during the day, three at night) to both groups over the circadian cycle (**Figure 1**). Before home cage monitoring, all CS/US mice showed normal fear acquisition, as demonstrated by increased percent freezing over time (**Figure 2B-C**), while female mice displayed decreased freezing responses compared to males (**Figure 1D**). On home cage recording day 1, in response to the CS in the home cage, the fear-conditioned pair-housed mice exhibited a strong CS-evoked freezing response compared to the CS-only pair-housed group (**Figure 2F and H**, also see video 1). These data demonstrate successful video based manual scoring of CS-evoked freezing behavior in the home cage of pair-housed mice CS-evoked freezing in the home cage of pair-housed mice across the light dark cycles: As shown in **Figure 2G and I**, and consistent with previous reports measuring CS evoked freezing across the light and dark cycle in novel arenas(Hartsock et al., 2024), we observed that within the home cage, pair-housed mice exhibited enhanced CS-evoked freezing during the light (inactive) period versus the dark (active) phase (**Figure 2G,I**). This increased percentage of CS-evoked home cage freezing persisted through day 6 of the study. After day 6, a CS-dependent home cage extinction effect was observed throughout the remainder of the home cage recordings (**Fig. 2F-J**). However, compared to males, female mice exhibited enhanced extinction, that plateaued on day 10 (**Figure 2J**). These data demonstrate successful circadian recordings of CS-evoked behavioral tracking, along with quantification of pair-housed CS-evoked freezing behavior, revealing sex-based differences in extinction learning rates.

**Fig 2.**
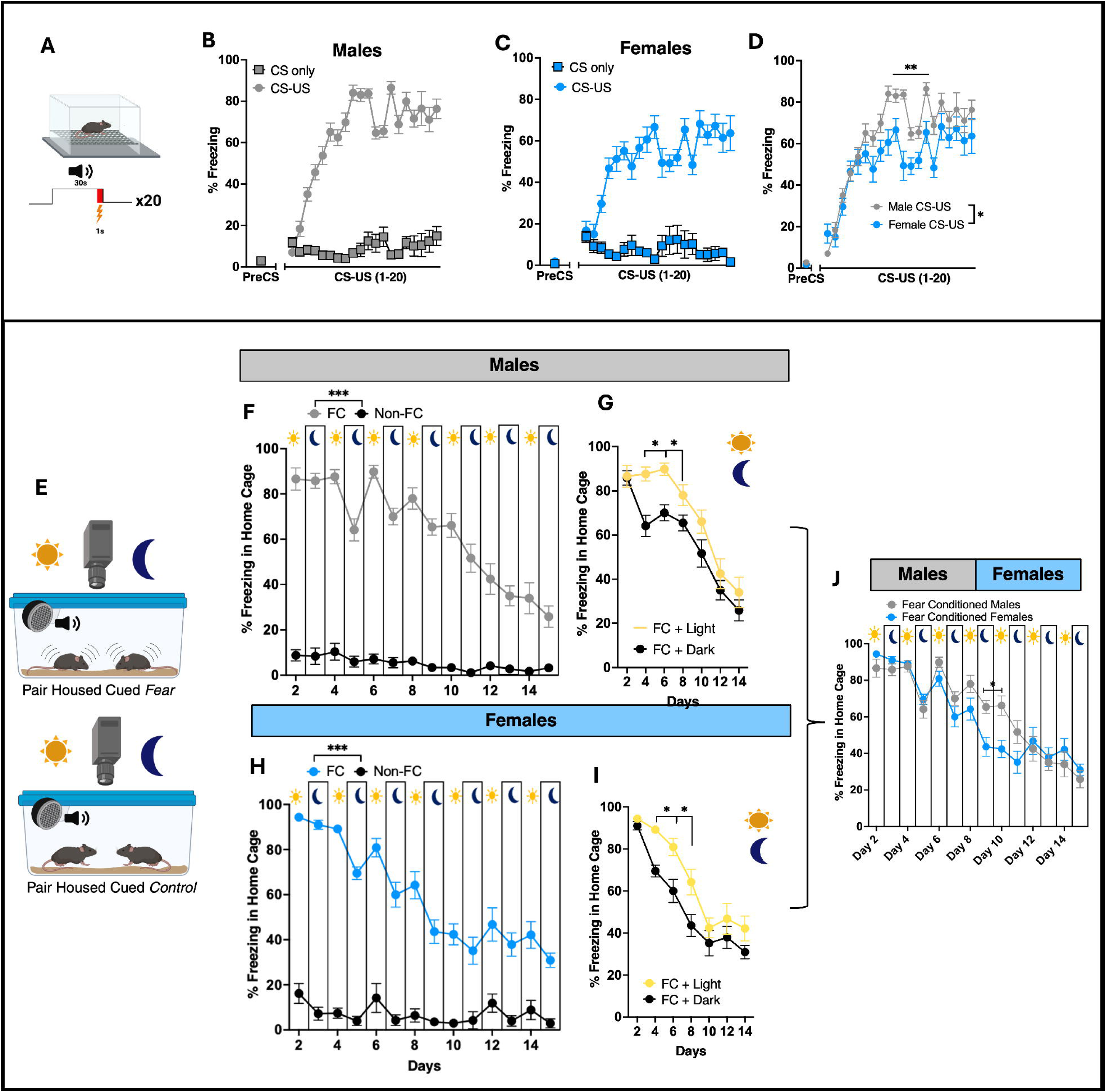
Assessment of Circadian Freezing Behavior Following Fear Conditioning in the Home Cage of pair-housed Mice. A. Schematic of fear conditioning protocol (20 CS/US pairings). B–C. Fear acquisition measured by percent freezing in male (n=16; B) and female (n=10; C) mice. D. Male mice show significantly greater freezing during conditioning compared to females (group effect: F(1,22) = 8.552, *p* < 0.01). E. Diagram of home-cage video monitoring system. F, H. Manual scoring of CS-evoked freezing in fear-conditioned (FC) vs. non-fear- conditioned (Non-FC) group-housed mice across 14 days (time: F(13,315) = 20.25, *p* < 0.0001; group: F(1,30) = 548.1, *p* < 0.0001; interaction: F(13,315) = 12.87, *p* < 0.0001). G, I. CS-evoked freezing in male (G) and female (I) mice across light/dark cycles (males: F(1,30) = 8.423, *p* = 0.0069; females: F(1,18) = 10.76, *p* = 0.0042). J. Sex differences in circadian CS-evoked freezing behavior (day–night × sex interaction: F(13,276) = 3.263, *p* = 0.0001).

### Deep learning-based pose estimation analysis for CS-evoked freezing analysis in pair- housed mice

Next, we aimed to validate our manual scoring of freezing behavior in the home cage using DeepLabCut (DLC), which allows for automated freezing analysis by labeling key points on the mouse’s body (24) (Mathis et al., 2018). All home cage video recordings were timestamped with the CS delivery from the ACD software and from these videos, a DLC algorithm was trained to track the movement of the nose, left ear, right ear, neck, body, and tail of each mouse within the home cage **(Video 2).** Once distance traveled and velocity were calculated using these points, we determined freezing based on low body point velocity. To track the freezing behavior of multiple mice in a dynamically changing home cage background, we converted the body point tracking data to freezing data **(Figure 3A-B).** After tracking with DLC, the raw tracking data was smoothed using the Simple Behavior Analysis (SimBA) software(Goodwin et al., 2024). As shown in **Figure 3C**, the automated tracking analysis was then validated through direct comparison with the manual scoring analysis, as illustrated in the representative video (see **Video 2)**. The freezing behavior of the mice in response to the CS was measured in 10-second intervals. Initially, freezing levels were comparable to baseline during the first 10-second interval, but they escalated to 90-100% during the second and third intervals. This data is consistent with the observed behaviors in the videos (**Video 2**), where mice dart to the corner of the cage at the beginning of the CS and freeze for the remainder of the time. In the non-fear-conditioned cage, the freezing response to the CS remained at baseline levels (**Figure 3D**), consistent with their behavior observed in **Video 2**. As shown in **Figures 3E-I**, a similar pattern between the automated and manual analysis of CS-evoked freezing was observed over 14 days, demonstrating the high performance of the automated tracking algorithm compared to manual scoring analysis. These results support the performance of our DLC trained algorithm and raw tracking data smoothed approach using SimBA and further demonstrate that this automated tracking method, is comparable to that of our manual scoring annotations and quantifications.

**Fig 3.**
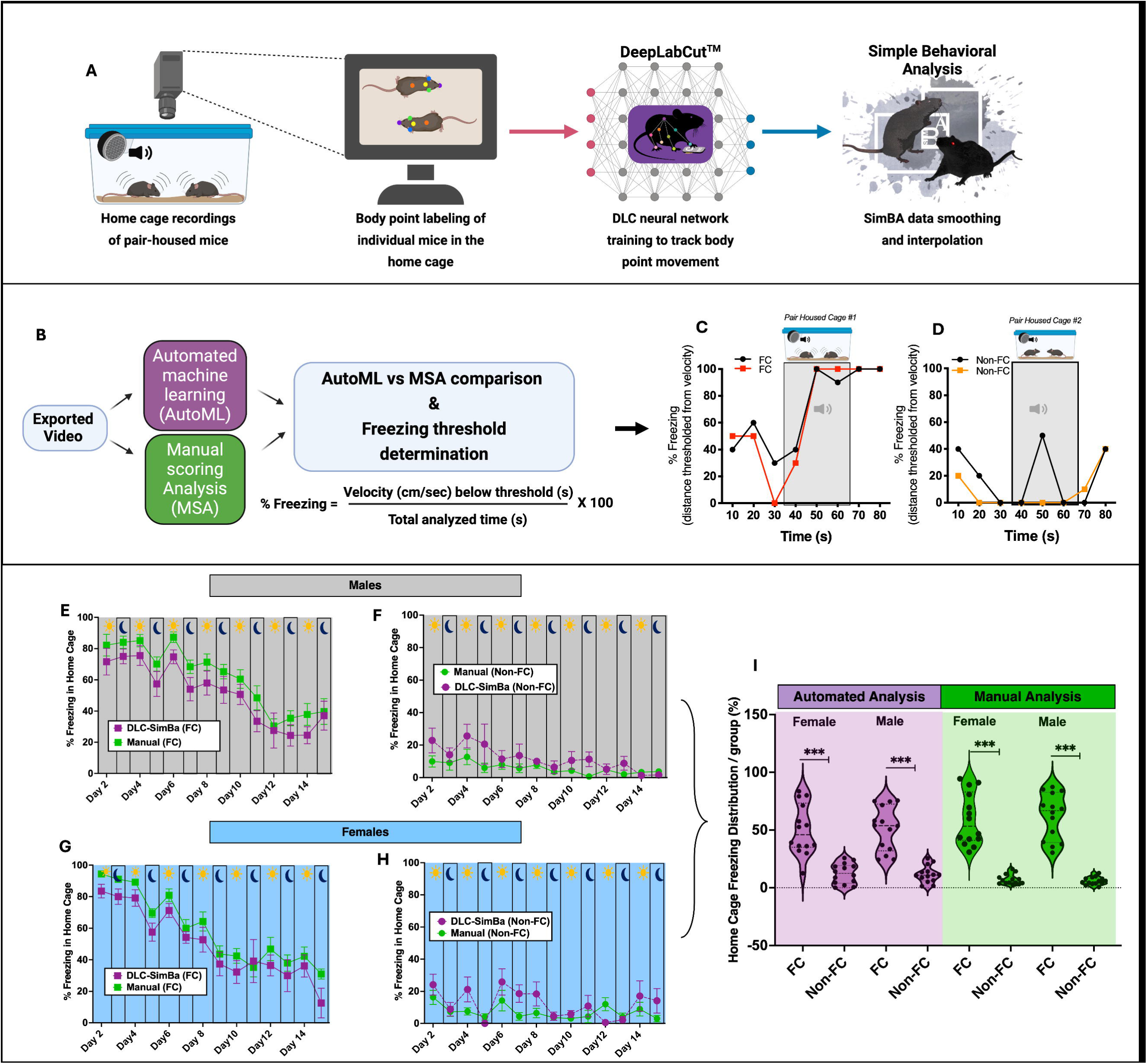
Machine learning-based tracking and analysis of CS-evoked freezing in pair- housed mice using DeepLabCut™ and SimBA™. A. Overview of experimental setup and automated tracking pipeline. B. Validation workflow for deep learning-based percent freezing detection. C–D. Automated analysis of thresholded freezing behavior (binned in 10-second intervals; see Video 2), converted to CS-evoked percent freezing in fear-conditioned (FC) and non-fear-conditioned (Non-FC) pair-housed mice (n=4 cages). E–H. Comparison of automated vs. manual scoring across 14 days in male Non-FC (E), male FC (F), female Non-FC (G), and female FC (H) groups. I. Violin plot showing the distribution of total CS-evoked percent freezing across all groups (n=8–12); treatment effect: F(7,104) = 37.24, *p* < 0.0001.

### Persistence of CS-evoked freezing in a novel context

At the end of the study, mice were removed from their home cage and tested for CS-evoked freezing in a novel context **(Figure 4A).** As shown in Figure 4B, both fear-conditioned male and female mice exhibited significantly higher CS-evoked freezing (fear expression or persistence test) in the novel context compared to CS-evoked freezing on Day 14 in the home cage **(Figure 4C).** Additionally, pre-CS freezing was low and similar between groups, indicating minimal contextual fear. These data suggest that despite CS-dependent extinction learning observed in the home cage during the latter part of the study (Days 8-14), enhanced cue-induced freezing persisted in a novel context, suggesting a potential renewal effect (Maren et al., 2013).

**Fig 4.**
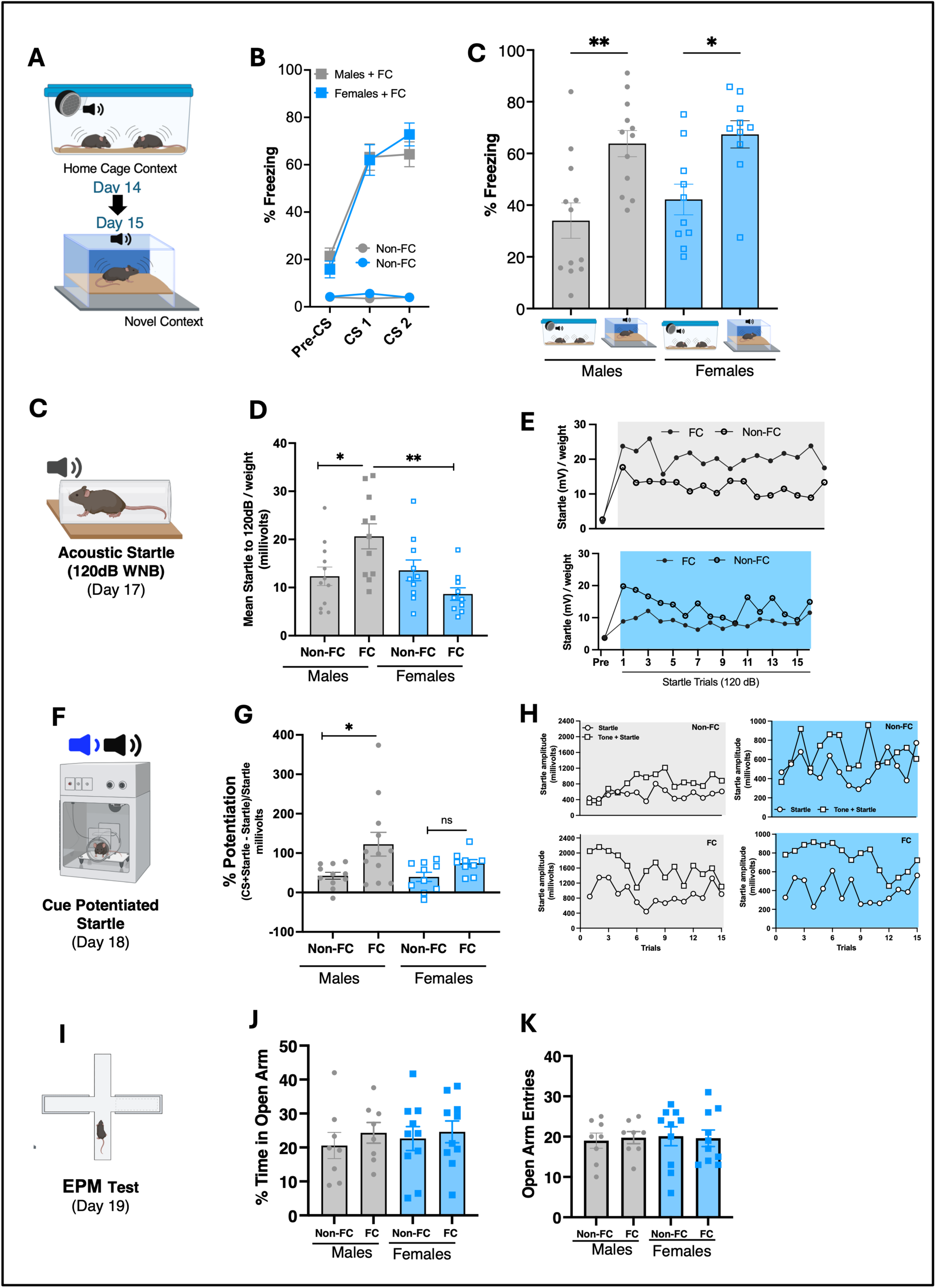
Sex-specific behavioral outcomes following fear conditioning (FC) and pseudo-randomized intermittent CS exposure in the home cage. (A) Experimental approach for assessing fear expression in a novel context (Day 15). (B) Percent (%) freezing across Pre-CS, CS1, and CS2 periods. FC groups (gray/blue) exhibited robust freezing, while Non-FC groups (light gray/light blue) did not. (C) FC-induced freezing in a novel context (Day 15) was greater in males than females (treatment effect: F(3,40) = 7.751, *p* = 0.0003). (D) FC males, but not females, show increased startle reactivity to a 120 dB white noise burst after 14 days of pseudorandomized CS exposure (treatment effect: F(3,40) = 17.03, *p* < 0.0001). (E) Startle leader trial traces for males (top) and females (bottom), showing trial-by-trial modulation by FC. (F) Schematic of fear-potentiated startle testing (Day 18), involving auditory CS presentations paired with startle stimuli. (G) FC males also show potentiated startle responses to the CS, not observed in females (treatment effect: F(3,38) = 3.632, *p* = 0.0213). (H) Startle traces from cue-potentiated startle test trials, separated by sex and condition. (I) Elevated Plus Maze (EPM) schematic for Day 19 avoidance / anxiety-like behavior testing. (J-K) % time spent in open arms (J) and number of open arm entries (K) were not significantly affected by FC in either sex.

### Enhanced baseline and fear potentiated startle (FPS) in males but not female mice

Following CS-dependent context testing, baseline startle reactivity, FPS, locomotor and avoidance-like behavioral testing **(Figure 4** **D,G)** was assessed. These data were analyzed and compared across sex and against the CS-only (auditory cue) control group. All mice were tested on separate days for both uncued (startle to 120dB white noise burst) and cued (fear-potentiated startle) reflexive startle responses. As shown in **Figure 4E**, the males but not females exhibited a significant enhanced uncued startle response. Similarly, when testing for FPS, the males but not females exhibited a significant enhanced CS-potentiated startle (**Figure 4G**) response. On the other hand, there were no differences between groups or sex when analyzed for locomotor, exploratory behaviors, as measured by EPM (**Figure 4I-K**). Taken together, these results demonstrate a sex-dependent enhancement in baseline startle and FPS responses, independent of locomotion or exploratory measures, in fear-conditioned mice exposed to repeated (pseudorandomized), intermittent auditory threat cues in a familiar (or “safe”) home cage environment.

## DISCUSSION

Here, we present a novel approach that integrates open-source software with home-cage video recording and deep learning–based pose estimation to enable pseudorandomized delivery of fear-conditioned stimuli (CS) to pair-housed mice in their home environment. The following are the key findings: (1) Demonstration of multi-day temporal video tracking of CS-evoked freezing behaviors in pair-housed mice across light-dark cycles in a home cage context; (2) Validation of a deep learning-based pose estimation algorithm for freezing behavior analysis in pair-housed mice; (3) The software was applied to generate a PTSD-like behavioral paradigm, revealing significant sex-dependent differences in CS-evoked diurnal freezing, startle reactivity, and fear renewal. Overall, these findings contribute to the advancement of integrative physiology and behavior analysis in home cage settings and may support future studies aimed at elucidating the mechanisms underlying heightened sensory-evoked physiological and autonomic reactivity in PTSD.

Enhanced physiological threat reactivity to trauma-related sensory triggers in non-threatening contexts (e.g., safe environments), and re-experiencing are core symptom clusters of PTSD criterion E, DSM-5(Association, 2013). PTSD-like animal models(Bale et al., 2019; Bowers and Ressler, 2015) are critical for investigating underlying mechanisms, including a more precise understanding of dysregulated sensory threat perception, such as heightened cardiovascular-autonomic responses to conditioned threat cues (Martin et al., 2024; Park et al., 2017; Seligowski et al., 2020). Using a Pavlovian-based aversive learning paradigm, a well-established preclinical approach for evaluating PTSD-related neurobiological mechanisms with strong relevance across species (Deslauriers et al., 2018; Dunsmoor et al., 2022), our first objective was to assess CS-evoked fear expression (i.e., freezing) in dyad (pair-housed) mice within a familiar home cage environment.

Fear in mice is typically assessed through behaviors like freezing and escape, as well as autonomic responses such as increased heart rate, posture changes, ultrasonic vocalizations, and piloerection(Fanselow, 1984; LeDoux, 2000). As demonstrated here, in fear-conditioned dyad mice, we observed the expected defensive behavioral pattern, a robust freezing response to the CS, on day 1 in a familiar environment (see supplemental videos). This increased CS-evoked fear expression in the home cage persisted up until the sixth day of the paradigm, while evidence of extinction learning (reduced CS-evoked freezing) emerged in subsequent days 8-14. Fear extinction is a form of new learning that results in the inhibition of conditioned fear, and deficits in fear extinction are a risk factor for anxiety disorders(Cain, 2023; Peters et al., 2009; Tronson et al., 2012; Velasco et al., 2023). To our knowledge, this study represents the first investigation to demonstrate and quantify dyadic CS-evoked freezing and extinction learning in pair-house mice in their home cages.

We next validated our manual scoring of freezing behavior using a multi-animal pose estimation DeepLabCut™ (DLC) approach (Chanthongdee et al., 2024; Goodwin et al., 2024; Mathis et al., 2018; Nath et al., 2019). Similar to Chanthongdee et al., we employed a combined DeepLabCut + Simple Behavior Analysis (Simba) workflow using a conditioned fear paradigm for freezing analysis. Using SimBA’s interpolation method we improved the mouse position tracking during frames with missing data caused by the mouse being out of view and poor tracking-related issues. In comparison to our manual scoring, our trained DeepLabCut + SimBA workflow demonstrated high inter-method reliability, including variations in circadian freezing over multiple days. Unlike our study, Chanthongdee et al. (2024) quantified additional fear behaviors (e.g., grooming, sniffing, rearing) along with conditioned freezing, but this study was designed for single rodent analysis within open arenas and operant chambers, rather than a multi-animal home cage environment as used here. In future studies, incorporating more complex fear behaviors (Chanthongdee et al., 2024) into a DLC-SimBA workflow, combined with the pair-housed home cage strategy used here, could enable a more comprehensive fear behavioral ethological analysis. Overall, these findings add to the growing field of deep learning-based pose estimation analysis in multiple mice, including light-dark phases (Gabriel et al., 2022; Kaul et al., 2024; Pereira et al., 2022) and considerations for temporal monitoring in ethologically relevant environments such as the home cage (Kahnau et al., 2023).

The effects of circadian rhythms on conditioned fear behaviors and the influence of different Zeitgeber phases (e.g., Z0 = light period, Z12 = dark period) are increasingly important yet mechanistically understudied (Albrecht and Stork, 2017; Bussi et al., 2024; Hartsock et al., 2024; Tsao et al., 2022). The behavioral recording system used here was designed to facilitate the recording and analysis of CS-evoked freezing during both light and dark phases. Similar to (Woodruff et al., 2018) we observed CS-evoked reductions in freezing behavior during the active (dark) phase compared to the non-active (light) phase, with more pronounced reductions in females than males. Behavioral conditioned freezing during the acquisition and extinction stages is influenced by circadian timing (light vs. dark phase) (Hartsock et al., 2024) (Albrecht and Stork, 2017; Chaudhury and Colwell, 2002). In the current study, all mice were fear-conditioned during the inactive light period (ZT6–ZT8), and therefore we cannot rule out the possibility that this timing influenced the reduced CS-evoked freezing observed at during the dark period (active phase). On the other hand, reduced CS-evoked freezing during the dark phase may reflect innate survival behaviors, as mice are less likely to freeze during their active phase (e.g., feeding, grooming) compared to the more vigilant light phase (Woodruff et al., 2018). While intriguing and relevant to the circadian biology of conditioned fear and PTSD, further studies are needed to explore these findings.

Based on our findings showing home cage-dependent extinction learning after the 14- day protocol, we next tested the effects of CS-evoked freezing outside the extinction context to assess the renewal of conditioned fear (Maren and Holmes, 2016). In Pavlovian rodent and human literature, it is well-established that fear extinction is context-specific (Maren, 2012; Maren et al., 2013). Supporting this concept, fear-conditioned mice that exhibited evidence of extinction learning (low CS-evoked freezing) in the home cage (context A), displayed high CS-evoked freezing when we tested them in a novel context (B), with no differences between males and females. These results suggest that mice undergoing extinction learning in a familiar home cage environment exhibit renewed or reactivated conditioned fear when the CS is delivered in a new environment (context B), further supporting the idea that extinction learning is context-dependent, with fear responses returning when contextual cues change(Bouton et al., 2006).

After fear renewal testing, to evaluate mechanisms of heightened arousal and avoidance-like behaviors, both groups were assessed for acoustic startle reflex (ACS), fear-potentiated startle (FPS), and elevated-plus maze (EPM). Fear-conditioned mice exposed to pseudorandomized CS’s in the home cage here, showed increased baseline startle and FPS compared to controls, independent of avoidance-like behavior. These findings align with previous rodent studies showing prior stress can enhance acoustic startle responses (Khan and Liberzon, 2004; Mancini et al., 2021; Sillivan et al., 2017; Torrisi et al., 2021). However, unlike these studies using innate physical stressors, our paradigm relied on repeated, pseudorandomized auditory cue reminders in the home cage, suggesting a distinct, yet undefined mechanism. Paradigms involving severe stressors that enhance startle and FPS often impact neuroendocrine, immune, and HPA axis function, contributing to behavioral changes(Ménard et al., 2017; Young et al., 2018). While we did not directly assess these systems, our conditioned fear model may produce enhanced startle through previously documented structural and synaptic plasticity in threat-related circuits (e.g., auditory cortex, amygdala, hippocampus) following fear conditioning and extinction (Gruene et al., 2016; Jr et al., 2015). Overall, these behavioral findings demonstrate that while CS-dependent extinction occurs in the home cage, conditioned fear persists in new environments (ie., renewal effect), and long-term CS-evoked behaviors in the home cage enhance baseline and fear-potentiated startle reactivity in males, but not females, independent of avoidance behaviors.

Sex differences in PTSD are well established, with women nearly twice as likely as men to develop the disorder in clinical populations (Olff, 2017; Pooley et al., 2018; Ressler et al., 2022). In our study, female mice showed reduced freezing during fear acquisition and home cage extinction compared to males, though no sex differences were observed in fear renewal or avoidance behaviors. Males, but not females displayed heightened startle and fear-potentiated startle responses after the two-week CS home cage protocol, which may be linked to greater initial fear acquisition and delayed extinction, though this requires further investigation.

Despite clinical findings showing women are twice as likely as men to develop PTSD, the rodent literature on sex differences in fear and stress responses (Bale & Epperson, 2015; Bangasser et al., 2018; Day & Stevenson, 2020), and as shown here do not always align with these clinical observations, making translation challenging. Variability in results are often attributed to factors such as ovarian hormone cycles, mouse strain, age, behavioral paradigm, and environmental conditions. These variables can impact stress and arousal systems (Bangasser et al., 2018), influencing behaviors such as freezing, darting, and startle reactivity (Olivera-Pasilio & Dabrowska, 2023; Mitchell et al., 2024). Overall, these findings are, to our knowledge, the first to report sex differences in a home cage extinction learning paradigm in pair-housed mice, while future studies are needed to clarify the neurobiological basis of these differences. These results may have important implications for understanding extinction learning across safe environments and emerging evidence of sex-specific fear synchronization behaviors (Velasco et al., 2019; Lebron-Milad et al., 2012; Ito et al., 2023).

In summary, this study introduces a novel PTSD-like behavioral paradigm supported by an open-source software platform, Adaptive Cue Delivery (ACD), that enables integrated video tracking, machine learning, and circadian analysis to deliver unpredictable pseudorandomized conditioned threat reminders in the home cage. This software and integrated behavioral approach may prove valuable in future studies investigating maladaptive fear regulation within ethologically relevant settings. Furthermore, integrating ACD with advanced imaging, circuit-level recordings, and closed-loop systems—such as telemetry-based methods (Turley et al., 2021), may advance translational research into the neural and physiological mechanisms underlying PTSD and its physical health comorbidities.

## Supporting information

https://gwu.box.com/s/wmfhizooznzmzrrwrys95t6mz9swqloi

https://gwu.box.com/s/kpoegjxkfk86atlkrxst2pvhnrbri6o0

https://gwu.box.com/s/c4z1lefi07qu23y839xjiyox9x13bfhl

https://gwu.box.com/s/5dumws1675hktpigozqazukarp081hng

https://gwu.box.com/s/kyspjqz4lhpr4yqrp9xx1fhf64tzm321

## Acknowledgments and Disclosures

This work was supported by Congressionally Directed Medical Research Programs (CDMRP) PR210574. PJM, HCS, ZY, JP conceived and designed the study. BT, AS ZY and HCS designed and tested the video analysis and CS scheduling software used in the study. HCS, ZY, PY performed the behavioral experiments. HCS performed the manual data scoring and analysis. ZY performed the automated DeepLabCut^TM^ and SimBa^TIM^ data scoring and analysis. Figures were created using Biorender. The authors declare no competing interests.

## Supplemental Files

S1: Table 1: Complete Statistical Output

S2. Table 2: **Timeline of behavioral testing and pseudorandom home cage CS exposure.** Mice were habituated in the home cage room before undergoing a two-day shock box habituation period (20 min and 40 min, respectively) followed by fear conditioning (Day 1).

Conditioned stimulus (CS) presentations were delivered in the home cage in a pseudorandomized on/off schedule across 7 test days (Days 3–24). Each test day consisted of a total of six CS trials at different timepoints, 3 CS across the light phase and 3 CS across the dark. Subsequent behavioral assays included fear expression testing in a novel shock box context (Day 24), Elevated Plus Maze (EPM, Day 25), pre-pulse inhibition (PPI, Day 26), and fear-potentiated startle (FPS, Day 27).

**S2: Methods:** Troubleshooting for DLC and SimBA deep learning algorithms

**S3 Video 1:** Representative video showing CS-evoked pair housed home cage freezing in the fear-conditioned (FC group - top frame), but not in the non-fear-conditioned (Non-FC group- bottom frame). Auditory cue plays at 13:52.

**S4 Video 2:** Representative video with body parts labeled using DeepLabCut^TM^, showing CS-evoked pair housed home cage freezing in the fear-conditioned (FC group - top frame), but not in the non-fear-conditioned (Non-FC group-bottom frame). Auditory cue plays at 10:18.

