## Supplementary material for "Tracking Conditioned Fear in Pair-Housed Mice Using Deep Learning and Real-Time Cue Delivery": https://gwu.box.com/s/kpoegjxkfk86atlkrxst2pvhnrbri6o0

### Supplemental machine learning algorithm methods and video recording set-up:

**Troubleshooting, Limitations, and Caveats of Automated Home Cage Freezing Analysis:** In our studies, the nose was generally the most reliable body part for detecting mouse movement using automated tracking. However, due to the constraints of our current recording system, particularly the limited field of view, the nose was frequently out of frame, resulting in it being the most common source of missing data. To address this, we used the ear as the primary tracking point. Minor movements involving only the nose were subsequently classified as freezing. This adjustment may have contributed to the trend toward reduced percent freezing observed in the DLC-shock group (see Fig. 3E–H). We anticipate that future improvements in recording conditions, including expanded fields of view or multi-angle setups, could enhance detection sensitivity and reduce reliance on manual correction.

**Supplemental Figure 1:** Video Recording System Set-up

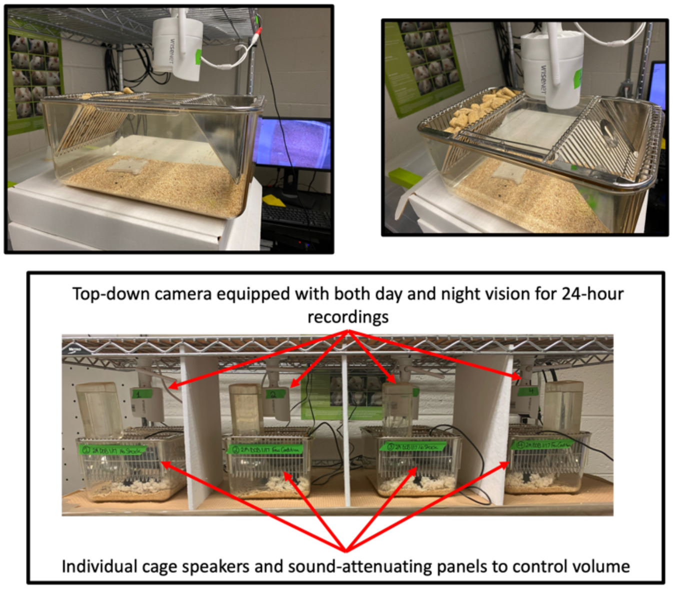


**Video Recording Equipment Setup (Figure 1, Inset):** The short working distance of the camera resulted in a limited recording field, and in-cage obstructions—such as food and water—likely contributed to the tracking limitations described above (see representative DLC inset and Supplementary Video). These constraints occasionally prevented the DeepLabCut (DLC) algorithm from reliably tracking certain body parts, particularly when mice were positioned in cage corners or rearing along the cage walls, leading to intermittent missing data points. To address this, we used SimBA's built-in interpolation method to estimate missing values. This was followed by manual review and correction to verify the accuracy of the interpolated data. Consequently, additional human analysis was required to support final comparative assessments between manual and automated scoring. Another limitation of the recording setup was frequent identity (ID) switches between the two mice in a pair-housed cage due to missing data, which complicated efforts to distinguish individuals. While this issue did not impact the current study, because both cage-mates received the same treatment (fear-conditioned or non-fear-conditioned) and displayed largely synchronized behavior, it limits the applicability of the approach for future studies involving non-synchronized or divergent individual behavior.

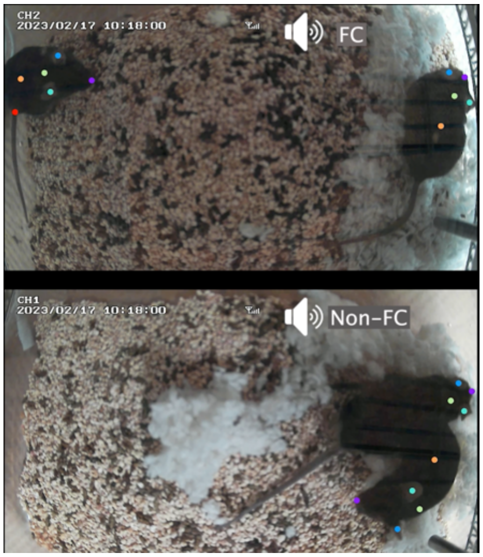
