## Supplementary material for "Tracking Conditioned Fear in Pair-Housed Mice Using Deep Learning and Real-Time Cue Delivery": https://gwu.box.com/s/c4z1lefi07qu23y839xjiyox9x13bfhl

### Supplemental 2: Table 2

| Timeline | Location | Days | Pseudorandom CS and Behavioral Test |
|---|---|---|---|
| | Home Cage Room | Habituation | Placed in cages 2pm |
| | Shock Box | 20m Habituation | 11am |
| | Shock Box | 40m Habituation | 10am |
| 1 | Shock Box | Fear Conditioning | 12pm |
| 3 | Homecage | Homecage CS Day 1 | 10:10, 13:52*, 17:34, 19:25, 2:49, 6:31 |
| 5 | Homecage | Homecage CS Day 2 | 10:22*, 17:46, 19:37, 4:52, 6:43, 8:34 |
| 7 | Homecage | Homecage CS Day 3 | 10:18*, 17:42, 21:24, 2:57. 4:48, 8:30 |
| 9 | Homecage | Homecage CS Day 4 | 12:12*, 18:04, 19:32, 21:00, 1:24, 11:40 |
| 11 | Homecage | Homecage CS Day 5 | 9:58*, 18:10, 0:19, 4:25, 6:28, 8:31 |
| 14 | Homecage | Homecage CS Day 6 | 10:56*, 13:44, 16:32, 22:08, 13:32, 5:08 |
| 16 | Homecage | Homecage CS Day 7 | 12:22*, 17:58, 23:34, 3:46, 6:34, 10:46 |
| 24 | Shock Box – novel context | Fear expression (2CS) | 12pm |
| 25 | Behavior room | EPM | 11am |
| 26 | Startle Box | PPI | 10am |
| 27 | Startle Box | FPS | 10am |
